# Photovoltaic stimulation efficiently evokes network-mediated activity of retinal ganglion cells

**DOI:** 10.1101/652404

**Authors:** Naïg A. L. Chenais, Marta J. I. Airaghi Leccardi, Diego Ghezzi

## Abstract

**Objective:** Photovoltaic retinal prostheses theoretically offer the possibility of standalone high-resolution electrical stimulation of the retina. However, in artificial vision, achieving locally selective epiretinal stimulation is particularly challenging, on the grounds of axonal activation and electrical cell coupling.

**Approach:** Here we show that electrical and photovoltaic stimulation of dystrophic retinal circuits with capacitive-like pulses leads to a greater efficiency for indirect network-mediated activation of retinal ganglion cells. In addition, a biophysical model of the inner retina stimulation is proposed to investigate the waveform and duration commitments in the genesis of indirect activity of retinal ganglion cells.

**Main results:** Both in-vitro and in-silico approaches suggest that the application of long voltage pulses or gradual voltage changes are more effective to sustainably activate the inner excitatory and inhibitory layers of the retina, thus leading to a reproducible indirect response. The involvement of the inhibitory feedback from amacrine cells in the forming of indirect patterns represents a novel biological tool to locally cluster the response of the retinal ganglion cells.

**Significance:** These results demonstrate that recruiting inner retina cells with epiretinal stimulation enables not only to bypass axonal stimulation but also to obtain a more focal activation thanks to the natural lateral inhibition. In this perspective, the use of capacitive-like waveforms generated by photovoltaic prostheses may allow improving the neural response resolution while standing high-frequency stimulation.

## 1. Introduction

Retinal dystrophies, such as age-related macular degeneration and retinitis pigmentosa, are ranked among the three leading causes of visual impairment worldwide (together with cataract and glaucoma) and are the primary cause of visual deficit in middle income and industrialized countries, with a prevalence above 15 % [1–3]. Though, inner and ganglion retinal neurons are known to be temporarily spared by the degeneration process and to be electrically excitable to convey artificial visual inputs to the lateral geniculate nucleus [4–7]. Several retinal prostheses have been developed in the past decade and demonstrated promising results to restore an elementary form of vision, including discrimination of high-contrast gratings, reading of large prints, and spatial orientation [7–13]. Nonetheless, current clinical implants provide limited visual acuity, and the sight quality is still far away from being adequate in daily life [14]. Spatial resolution keeps being one of the biggest challenges (together with the wideness of the visual field) to achieve a valuable artificial vision restoration. To overcome those challenges, we have designed a wireless epiretinal prosthesis (POLYRETINA) able to restore a theoretical visual acuity of 20/600 and a visual field of 46 degrees, thanks to miniaturized photovoltaic pixels made of organic semiconductors [15].

The stimulation resolution in epiretinal configuration could be improved in various complementary ways: minimizing the spread of the electric field generated by single electrodes to avoid stimulation crosstalk [15, 16], reducing the electrode to retinal ganglion cell (RGC) layer distance to enhance stimulation specificity [17–19], increasing the charge injection capacity of microelectrodes [20–22], identifying the stimulation protocol able to selectively activate RGCs nearest the electrode [23, 24], or selecting the electrode array features curtailing the activation of RGCs distal axon segments [25, 26]. Indeed, the poor visual acuity reported in patients is often associated with the perception of amplitude-dependent elongated phosphenes [27, 28], attributed to the local hyperpolarization of axons of passage located between the epiretinal array and the RGC layer [7,25,26]. This local depolarization can be antidromically propagated until the ganglion cell soma located further away from the electrode, leading to diffuse ovoid activation of the retinal map, thus making it difficult for an implanted patient to perceive complex shapes [29, 30]. Though the axon initial segment is the compartment exhibiting the lowest activation threshold [29], in practice, its activation threshold is barely discriminable from the ones of more distal axonal segments [26], making spatially selective stimulation of RGCs a challenge.

To overcome this issue, a large share of effort has been directed towards the understanding of indirect stimulation of RGCs [31, 32], which arises from the activation of presynaptic neurons, such as bipolar cells (BCs) and/or photoreceptors (PRs), which in turn lead to a secondary network-mediated excitation of RGCs [31,33–37]. In general, subretinal prostheses are considered to be more efficient in targeting those presynaptic neurons, due to their location [30, 31]. However, the long-term efficacy of subretinal prostheses could be limited by other aspects, such as a more complex surgical approach, the limited possibility of replacement, or the presence of a glial seal upon PR degeneration. Nonetheless, although epiretinal arrays deliver stimuli from the RGC side, they have also shown the ability to elicit indirect stimulation [31, 33], which makes the epiretinal configuration particularly advantageous to bypass the aforementioned problems while providing a network integrated form of stimulation. Although no clear discrimination can be made between direct and indirect activation thresholds for epiretinal configuration [15, 31], it has been established that pulses of short duration (shorter than 0.7 ms) preferentially directly activate RGCs (e.g. by activation of their axons), generating a single action potential with high temporal precision [34, 35]. On the contrary, pulses of longer duration also lead to an additional indirect excitation of RGCs [31,33,35,36,38]. Indirect firing patterns have been reported to occur within 10 to 70 ms after the stimulus delivery, with a great variation between experimental designs [31–38]. Despite this apparent lack of temporal precision, it has been demonstrated that electrically induced indirect responses of RGCs closely matches with the spatiotemporal complex light-evoked responses [39]. Both the Argus^®^ II retinal prosthesis and the Alpha-IMS implant, the two systems currently approved by regulatory bodies, focus on temporal precision and respectively use pulses of 0.45 ms [8] and 1 ms [9]. However, a recent study pointed out the relationship between the inner retina activation elicited by long pulses and the achieved spatial resolution both in-vitro and in patients implanted with the Argus^®^ II device [24]. All these pieces of evidence suggest a clinical relevance of indirect activity to improve the resolution of retinal responses upon electrical stimulation. However, little is known about the exact mechanisms behind the epiretinal indirect activation of RGCs. Converging evidence suggests the activation of the inner retina layer as the main factor causing the biphasic spiking pattern of RGCs [15,31,36,40,41]. Because of their elongated shapes, spared PRs are as well susceptible to be activated by extracellular voltage gradients around the stimulation site. Moreover, taking into consideration the milliseconds latency between direct and indirect activities, together with the oscillatory spiking pattern reported notably in brisk-transient RGCs [38], it is likely that the generation of a network-mediated response involves the coordinate activation of several cell types, including the retinal inhibitory network.

Our previous work with POLYRETINA has shown that light sensitivity could be restored in retinal degeneration 10 (rd10) mice retinas at advanced stages of degeneration thanks to photovoltaic stimulation [15]. Consistently with other groups’ findings, we reported both direct and indirect activation of targeted RGCs, upon delivery of 10-ms light pulses. Although pulses with a rectangular shape are conventionally used in retinal prostheses, our photovoltaic interfaces intrinsically generate non-rectangular capacitive-like photocurrent and photovoltage pulses [15]. Growing insights suggest that non-rectangular pulses can elicit stronger network-mediated RGC activity than rectangular ones; in fact, stimulus shapes with low charge increase rates are able to elicit a stronger activation of the presynaptic neurons, leading to increased RGC indirect firing activity. Conversely, low-pass filtering inner retinal cells are less activated by high-frequency signals such as rectangular ones, more favourable to the direct depolarization of RGCs [42, 43].

Given the importance of selective network-mediated stimulation, we explored whether and how our photovoltaic approach alters the response pattern of RGCs by favourably eliciting network mediated activity instead of direct RGC depolarization. Our findings commit to being generalised to improve the spatial selectivity of epiretinal prostheses.

## 2. Methods

### 2.1 Electrophysiology

Experiments were conducted according to the ethical authorization GE3717 approved by the Département de l’emploi, des affaires sociales et de la santé (DEAS), Direction générale de la santé of the République et Canton de Genève, Switzerland. RGC activity was recorded from rd10 mice at post-natal days (P) 144 ± 18.5 (mean ± s.d). Eyes were enucleated from euthanized mice (sodium pentobarbital, 150 mg kg^−1^) and dissected in carboxygenated (95% O_2_ and 5% CO_2_) Ames’ medium (A1420, Sigma-Aldrich). Retinas were mounted ganglion cell down and maintained in contact with the substrate using a 1 mm nylon mesh. Retinas were continuously superfused with carboxygenated Ames’ medium at 32°C and maintained under dim red light during all the experiments.

Photovoltaic stimulation was carried out with the central part of the POLYRETINA photovoltaic array, consisting of 80-μm diameter electrodes distributed with a 150-μm pitch [15]. Electrical stimulation was carried out with a custom-made microelectrode array (MEA), consisting in a grid of 16 × 16 (256) titanium electrodes (80-μm diameter) distributed with a 150-μm pitch. Retina explants were illuminated with an inverted microscope (Ti-E, Nikon Instruments) and a LED illuminator (Spectra X, emission filter 560/32 nm, Lumencor). The microscope was equipped with a dichroic filter (FF875-Di01–25 × 36, Semrock) and 4x / 10x / 20x objectives (diameter of the illumination spot 5.5, 2.2, and 1.1 mm respectively; CFI Plan Apochromat Lambda). The stimulation protocol consisted of a repetition of 10 pulses at 1 Hz for each condition. Irradiance, stimulus shape, and pulse duration were increased sequentially up to the condition eliciting the highest RGC activity.

In experiments involving photovoltaic and electrical stimulation, the activity of RGCs was recorded extracellularly with a sharp metal electrode (PTM23BO5KT, World Precision Instruments), amplified (Model 3000, A-M System), filtered (300 - 3000 Hz), and digitalized at 30 kHz (Micro1401–3, CED Ltd.). Spike detection and sorting were performed by threshold detection using a MATLAB-based algorithm (Wave_clus47 [44]); results were further processed with MATLAB (Mathworks). An exclusion period of ± 1 ms around light onset and offset was applied and spikes detected in the first 10 ms after light onset were manually verified to ensure a proper artefact rejection. Spikes raster from 10 consecutive sweeps were averaged and discretize to compute 10-ms bins PSTHs. Spikes were classified into short, medium, and long latency (respectively SL, ML, and LL) according to their timing after light onset, as in a previously described procedure [15]. The electrical receptive fields from individual RGCs upon electrical stimulation were centred on the electrode eliciting the maximal ML activity under 50-ms stimulation and normalized according to the ML firing rate achieved. The electrical receptive field (eRF) diameters were calculated as the full width at median response amplitude of experimental firing rates fitted distributions. For the assessment of the natural light responsivity, retinas from rd10 mice at various ages were mounted on filter paper and placed ganglion cell down on a transparent MEA with 256 electrodes (256MEA200/30iR-ITO, Multichannel Systems). The voltages of the 256 recording electrodes were amplified, filtered (300 – 3000 Hz), and digitalized with a 10 kHz sampling frequency (USB-MEA256-System, Multichannel Systems). Spike sorting from recordings were performed with MC_rack software (V 4.6.2, Multichannel systems); results were further processed with Neuroexplorer (v4, Neuronexus) and MATLAB.

### 2.2 Computational model

Biophysical retinal layers were modelled using Python-based NEST 2.14.0 Simulator tool [45]. A grid of 10 × 10 unspecific RGCs was modelled as Hodgkin–Huxley neurons, connected with gap junctions and placed 10 µm above an 80-µm diameter epiretinal electrode. Inner retinal cells were modelled as non-spiking integrating neurons, whose parameters are summarised in Table 1.

**Table 1.** Retinal layer’s parameters

| Neurons layer | RGC | AC | BC | HC |
| --- | --- | --- | --- | --- |
| Ratio to RGC | 1 | 5 | 4 | 1 |
| Distance to electrode | 10 µm | 30 µm | 49 µm | 64 µm |
| Cell diameter | 10 µm | 5 µm | 5 µm | 10 µm |
| Threshold potential | -55 µV | - | - | - |
| Resting potential | -70 µV | -70 µV | -70 µV | -70 µV |

The layers filters’ time constants were calculated as in equation (1).

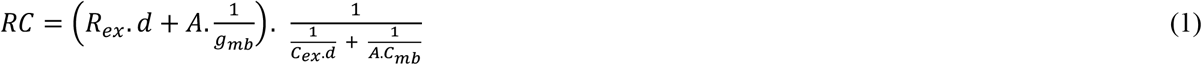

where *d* represents the cell distance to the electrode surface and *A* the area of bilayer membrane in the ascending column between the targeted cell and the electrode, as in equation (2):

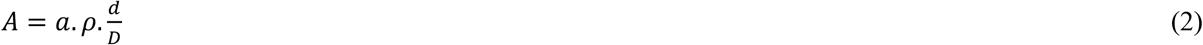

in which ρ represents the horizontal cell density in a theoretical receptive field and *D* an average retinal cell diameter. Parameters values are provided in table 2.

**Table 2.** Biophysical parameters of the retinal network

| $R_{ex}$ | $C_{ex}$ | $C_{mb}$ | $g_{mb}$ | $a$ | $\rho$ | $D$ |
| --- | --- | --- | --- | --- | --- | --- |

| **Extracellular resistivity** | **Extracellular capacitance** | **Passive membrane capacitance** | **Bilayer membrane conductance** | **Single cell surface** | **Cell density in the receptive field** | **Average retinal cell diameter** |
| --- | --- | --- | --- | --- | --- | --- |
| [$\Omega\text{m}^{-1}$] | [$\text{Fm}^{-1}$] | [$\text{Fm}^{-2}$] | [$\mu\Omega^{-1}\text{m}^{-2}$] | [$\mu\text{m}^2$] | [-] | [$\mu\text{m}$] |
| 10 | 0.1 | 0.01 | 5 | 300,000 | 42.8 | 7 |

The voltage density probability around and above the epiretinal electrode was simulated with a finite element analysis (FEA) method (COMSOL Multiphysics® v. 5.2.), with a stationary electric current study. The ground was situated at the bath top and lateral walls, located respectively 2 mm and 1 mm away from the studied pixel. The retinal tissue was placed from 10 to 110 µm above the electrode surface. The distribution of the voltage densities in the retina was evaluated every 5 µm from the retinal surface. For each material, the conductivity (S m^−1^) and relative permittivity values were set to: titanium (2.6 × 10^6^ / 1), P3HT:PCBM (0.1 / 3.4), PEDOT:PSS (30 / 3), Saline (1 / 80), PDMS (2 × 10^−14^ / 2.75), and retinal tissue (0.1 / 0.01). Prosthetic voltage density for artificial neurons stimulation was spatially approximated as a Gaussian probability distribution and temporally fitted as a first-degree exponential peaking at the FEA output values. The membrane potential of the cells located in the centre of the stimulated electrode, the corresponding RGC PSTH, and the two-dimensional firing activity was directly assessed using the multimeter tool. The AC activity level was evaluated with respect to the minimum AC membrane potential allowing network-mediated activity (−21.53 mV). The spatial extent of the RGC activity was calculated as the full width at half maximum of the RGC population activity gaussian fit. The spatial extent of the AC activity was assessed as the full width to AC membrane potential activation threshold of the AC membrane potential fitted distribution.

## 3. Results

### 3.1 Photovoltaic stimulation of the inner retinal network

All the experiments have been performed with explanted retinas from rd10 mice, which is an established model for retinitis pigmentosa [46–49]. However, in order to formally exclude any intrinsic light responsivity from possible surviving PRs, we first assessed the time course of the light responsivity decay at the wavelength and irradiances used for prosthetic stimulation with POLYRETINA, namely full-field light pulses of 10 ms at 560 nm, with irradiances ranging from 0.39 to 27.2 mW mm^-2^ (**Fig. 1a**). The relative percental increase or decrease in firing rate during a 180-ms time window after light onset (with respect to a 100-ms time window before the light onset) has been measured to determine a light responsivity index accounting for both ON and OFF transient and sustained responses (*N* = 4 retinas per timepoint, all RGCs recorded for each retina have been averaged). Rapidly after the formation of functional PRs at P16, a rapid decay in the light responsivity index has been observed up to P60, where it reaches its minimum value and remains constant around a baseline value of 0 regardless of the irradiance used (**Fig. 1b**). The time course of this loss of light sensitivity is in line with the reported anatomical changes in the outer nuclear layer of rd10 retinas [42, 43]. In young retinas (from P16 to P45), mostly transient response patterns could be detected (**Fig. 1a**), with a mean (± s.d.) latency of 69.8 ± 10 ms. In such light-sensitive retinas, green light responsivity increases with stimulus intensity up to 1.3 mW mm^-2^; after which irradiance increase weakens the average retina’s response, most probably because of M-cones saturation and eventually bleaching for the highest values of intensities tested (higher than 9 mW mm^-2^), as visible at the P16 time point. In retinas over P60, no significant light responding RGC could be recorded. Contrarily, the vast majority of RGCs from retinas explanted at those advanced stages of degeneration exhibits a robust light-independent spontaneous activity pattern with a peak frequency of about 10 Hz, in line with previous reports [48–50]. The more advanced the degeneration process of the recorded retinas, the higher the number of cells presenting strong spontaneous activity could be observed, with a peak around P100. To ensure a proper exclusion of intrinsic light-responses together with a proper detection of functional RGCs, further experiments have been performed on rd10 retinas at late stages of degeneration (i.e. after P120).

**Figure 1.**
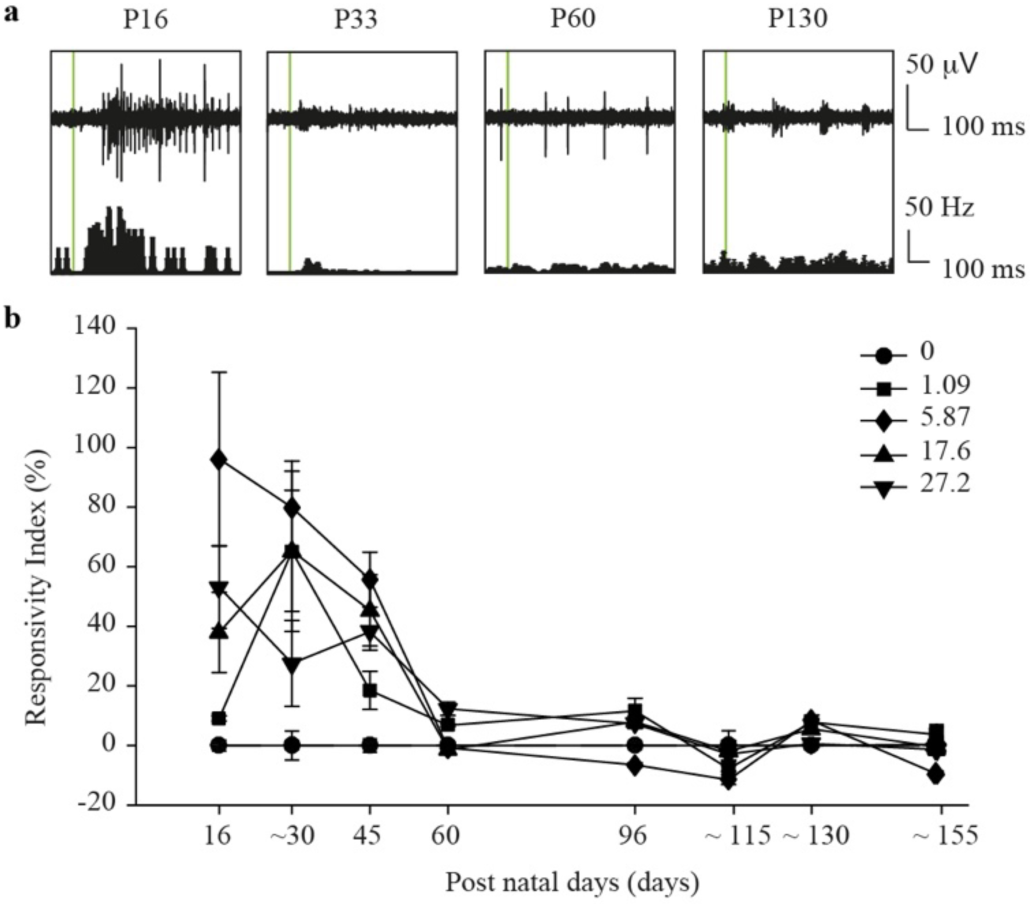
Light responsivity decay in rd10 retinas over the degeneration process. **a** Representative spiking activity from rd10 RGCs explanted at P16, P33, P60, and P130 in response to a light pulse (10 ms, 560 nm, 1.09 mW mm^-2^). **b** Mean (± s.e.m, *N* = 4 retinas) light responsivity index upon illumination (10 ms, 560 nm) of rd10 retinas recorded at P16, [P28-P38], P45, P60, P96, [P109-P116], [P128-P133], and [P153-P156]. For each RGC 10 sweeps have been averaged. Data have been shown only for five irradiances: 0, 1.09, 5.86, 17.6, and 27.2 mW mm^2^.

Then, we investigated whether with the photovoltaic approach the duration of the light pulse alters the response pattern of RGCs, similar to what has been reported for electrical stimuli [31, 32]. Retinas have been explanted over the POLYRETINA prosthesis with 80-µm photovoltaic pixels in epiretinal configuration and the activity of RGCs (*n* = 16 from *N* = 9 retinas) has been recorded upon light pulses of 10, 20, 50, and 100 ms, with irradiances ranging from 1.09 to 91.59 mW mm^-2^. Consistently to other reports [8,9,14,15], extracellular recordings under photovoltaic stimulation reveal a spiking pattern of RGCs made of two to three waves of activation, referred as SL, ML, and LL activities. Direct activity (SL) is theoretically elicited within a time window of few milliseconds from the pulse onset. As we have previously documented with POLYRETINA, such direct activation can be evoked from a low irradiance threshold (47.35 µW mm^-2^) [15]. Within the tested irradiance range (from 1.09 to 91.59 mW mm^-2^) and pulse durations (from 10 to 100 ms), SL spikes are elicited at an average (± s.e.m.) stable frequency of 36.9 ± 8.25 Hz (**Fig. 2d**, top). Instead, indirect (ML and LL) activity, originating from the activation of the upstream retinal network, strongly depends on the light exposure, both on irradiance and on pulse duration (**Fig. 2b-d**). As previously reported [15, 31], irradiance thresholds for direct and indirect activities are barely discriminable, and ML activity could be recorded from the lowest irradiance tested (1.09 mW mm^-2^). LL activity requires higher exposure to appear; it has been detected from an irradiance of 11.68 mW mm^-2^, for the longest pulses only (**Fig. 2b**, red arrow), or higher for shorter pulse duration. The independent increase of either irradiance or pulse duration allows the strengthening of the indirect responses (ML and LL). Over-threshold indirect activity (ML) rose up to 68 ± 8.3 % (mean ± s.e.m) of its initial values with an exposure increase from 1.09 to 91.59 mW mm^-2^ (average of all pulse durations tested). Besides, over-threshold indirect activity rose by 44 ± 7.4 % (mean ± s.e.m) when lengthening the stimulation time from 10 to 100 ms (average of all irradiances tested). The highest indirect (ML) activity is recorded at maximal irradiance and pulse duration conditions (100 ms, 91.59 mW mm^-2^) and could go up until 135.83 ± 8.25 Hz (mean ± s.e.m.). Noteworthy, not all the recorded RGCs showed a third wave of activity (LL), independently of the animal age and retina quadrant recorded. It is also worth to mention that RGC stimulation, if it does not completely abolish cells’ spontaneous activity, disrupts the excitatory oscillation during the evoked spiking pattern (**Fig. 2a**).

**Figure 2.**
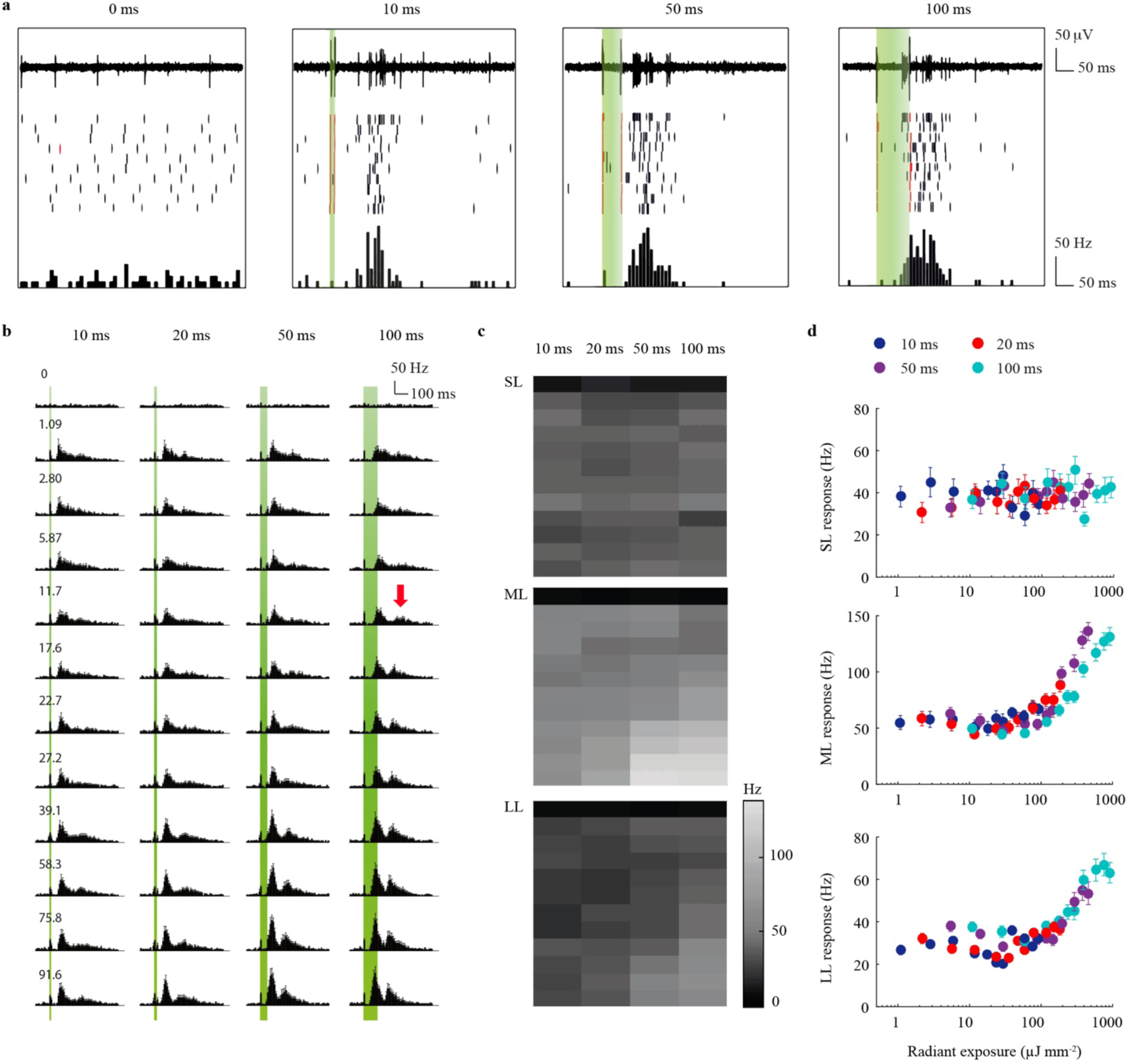
Photovoltaic stimulation elicits exposure-related activity in rd10 retinas. **a** Representative extracellular spiking activity of a single RGC in response to 0, 10, 50, and 100 ms photovoltaic stimulation with POLYRETINA at 2.80 mW mm^-2^. The top row shows the electrophysiological recordings upon illumination; the middle row the raster plot of 10 consecutive sweeps; the bottom row the PSTH (bin size 10 ms). The red raster lines correspond to the detection of the stimulus artefacts. **b** Mean (± s.e.m) PSTH (bin size 10 ms) of RGC activity upon 10, 20, 50 and 100 ms photovoltaic stimulation (*n* = 16, for each RGC 10 sweeps have been averaged). Light pulses with increasing irradiances have been delivered successively at 1.09, 2.80, 5.87, 11.68, 17.60, 22.76, 27.17, 39.06, 58.32, 75.80, and 91.59 mW mm^-2^. The red arrow indicates the onset of LL spikes. In **a** and **b**, the green bars correspond to the duration of the light pulse. **c** Heatmaps of SL (top), ML (middle), and LL (bottom) mean firing rates upon 10, 20, 50, and 100 ms photovoltaic stimulation. The irradiance increases from the top row towards the bottom. **d** Mean (± s.e.m) firing rates of SL (top), ML (middle), and LL (bottom) responses, computed for all the tested exposures (*n* = 16, for each RGC 10 sweeps have been averaged) and plotted as function of the radiant exposure (μJ mm^-2^), obtained by multiplying the irradiance (μW mm^-2^) per the pulse duration (s). Firing rates corresponding to 10, 20, 50, and 100 ms are respectively plotted in blue, red, violet and cyan.

### 3.2 Photovoltaic versus rectangular stimulation of the inner retinal network

Second, we investigated whether the specific pulse shape of the photovoltage delivered by POLYRETINA had an impact on its ability to activate the inner retinal network. Retinas have been explanted in epiretinal configuration over a custom-made MEA with 256 titanium electrodes of 80-µm in diameter and electrically stimulated with voltage-controlled pulses; the evoked spiking activity of RGCs has been recorded as before (**Fig. 3a**). The mean photovoltage profile (red curve in **Fig. 3b**, data profile obtained from [15]) generated by the photovoltaic pixels (light pulses of 10 ms and 0.94 mW mm^−2^, leading to a peak voltage of 179 mV) has been scaled to generate capacitive-like voltage pulses of various peak amplitudes. In a first subset of retinas, both 10-ms anodic and cathodic capacitive-like profiles of increasing peak voltages (8.95, 17.9, 35.8, 179, 368, and 895 mV) were randomly injected through the MEA electrode closest to the monitored RGC (*n* = 10 cells from *N* = 4 retinas). In agreement with the hypothesis that network-mediated activity is elicited by the direct transmembrane depolarization of the inner retinal cells, the threshold for network-mediated activity is lower in case of the cathodic profile injection with respect to the anodic one (−8.95 mV vs. +17.9 mV, respectively) for ML activity (**Fig. 3c**, top). The LL activation thresholds did not show any clear trend due to LL high intrinsic cell-to-cell variability (**Fig. 3c**, bottom). In a second subset of retinas (*n* = 13 cells from *N* = 11 retinas), a wider range of cathodic capacitive-like pulses has been successively injected (from −8.95 to −1790 mV). No statistical difference has been found between RGC activity elicited by photovoltaic stimulation (1.09 mW mm^−2^) and the corresponding capacitive-like voltage profile (**Fig. 3d**; SL: p = 0.17; ML: p = 0.16; LL: p = 0.98; t-test).

**Figure 3.**
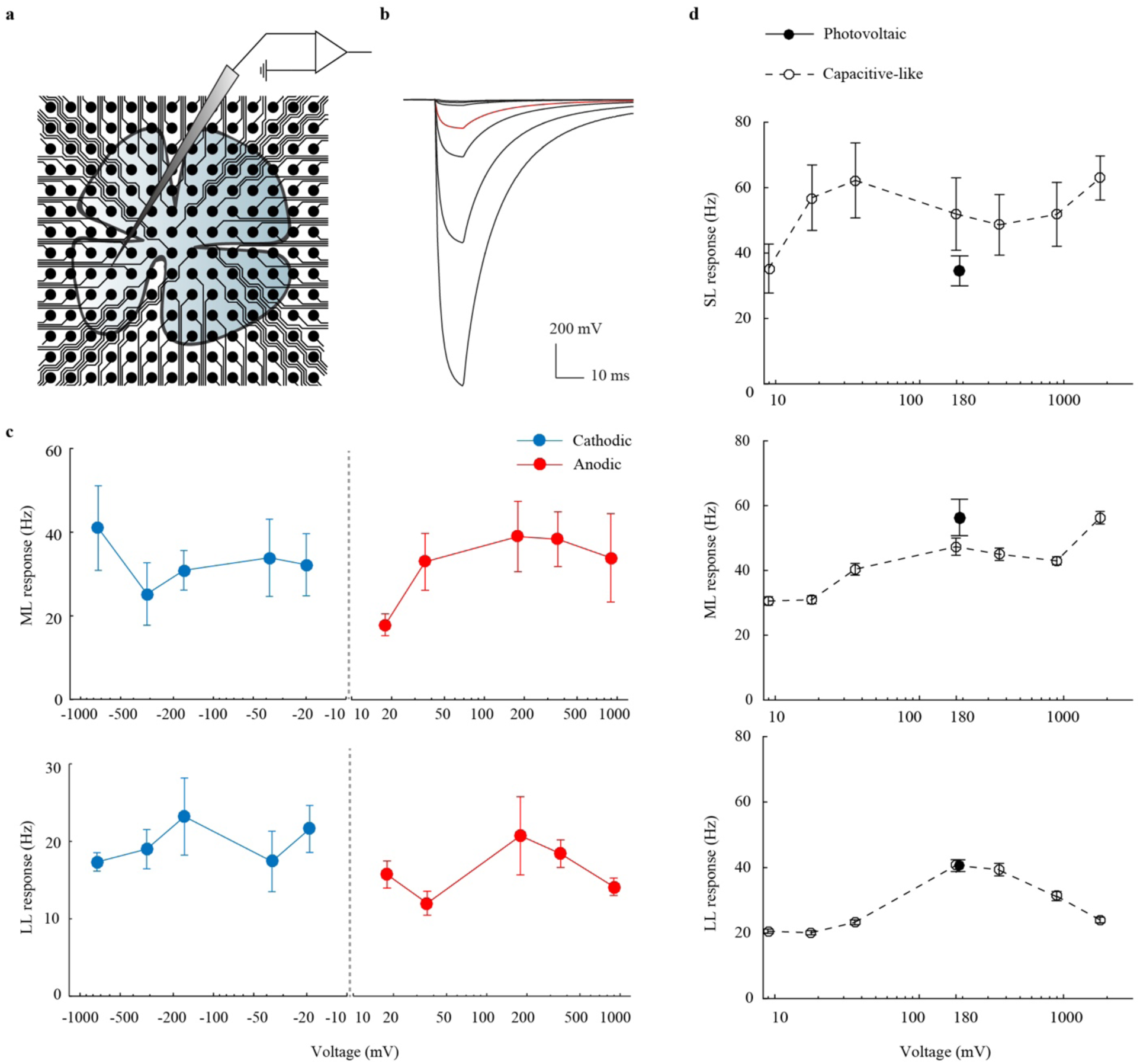
Electrical stimulation of rd10 retinas with capacitive-like voltage pulses. **a** Sketch of the stimulating and recording set-up. **b** Capacitive-like waveforms used for electrical stimulation. The red trace corresponds to the mean photovoltage generated by a photovoltaic pixel of POLYRETINA upon a light pulse of 0.94 mW mm^-2^ at 565 nm for 10 ms (data profile obtained from[15]) with a peak amplitude of 179 mV. The black traces show the capacitive-like voltage profiles obtained by scaling the peak amplitude to peak voltages of 8.95, 17.9, 35.8, 368, 895 and 1790 mV. **c** Mean (± s.e.m) ML and LL spiking activities in response to anodic (red) or cathodic (blue) electrical stimulation with capacitive-like voltage pulses (*n* = 10, for each RGC 10 sweeps have been averaged). **d** Mean (± s.e.m) SL, ML and LL activities upon photovoltaic stimulation (10 ms, 1.09 mW mm^-2^; *n* = 16, for each RGC 10 sweeps have been averaged) and electrical stimulation with capacitive-like pulses (*n* = 13, for each RGC 10 sweeps have been averaged).

Next, we compared the spiking activity of RGCs (*n* = 13 cells from *N* = 11 retinas) evoked by rectangular voltage pulses to the one evoked by capacitive-like pulses. Each RGC has been successively stimulated with rectangular pulses and capacitive-like pulses of identical peak values and of three different durations: 10, 50, and 100 ms. As before, to obtain capacitive-like pulses of 50 and 100 ms duration, the mean photovoltage profile generated by the photovoltaic pixels with pulses of 50 and 100 ms (0.94 mW mm^−2^, data profiles from [15]) have been scaled to generate capacitive-like voltage pulses of various peak amplitudes. Altogether, in agreement with the previous set of experiments, direct activity (SL) could be elicited from an average (± s.e.m.) voltage of 151 ± 12.9 mV and saturated for stimulation amplitudes higher than 358 mV (**Fig.4**). Direct activation threshold was not significantly different between capacitive-like and rectangular pulses (p = 0.37, t-test). On the contrary, indirect activity (ML and LL) is voltage and duration dependent. For identical pulse durations, the mean (± s.e.m) voltage thresholds obtained with capacitive-like pulses (132 ± 49.5 mV for 10-ms pulses; 149 ± 35.5 mV for 50-ms pulses; 170 ± 46.1 mV for 100-ms pulses) were significantly lower (p < 0.01 for all pulse durations, t-test) than the thresholds obtained with rectangular pulses (695 ± 124 mV for 10-11 ms pulses; 559 ± 128 mV for 50-ms pulses; 577 ± 128 mV for 100-ms pulses). Capacitive-like pulses elicited a mean (± s.e.m) direct activity of 47.5 ± 2.72 Hz, and rectangular pulses elicited a mean (± s.e.m) direct activity of 29.3 ± 1.40. The mean (± s.e.m) ML and LL spiking activities elicited with capacitive-like profile rose respectively to 41.9 ± 2.94 Hz and 28.5 ± 3.44 Hz, and to 31.4 ± 2.78 Hz and 26.3 ± 2.29 Hz with rectangular ones (**Fig.4c**). The mean (± s.e.m) cells’ indirect activity was inflated up to 16 ± 3.9 % of its initial values by increasing the applied peak voltage from the minimal condition eliciting indirect spikes to the maximal tested condition (from 358 mV to 1790 mV, average of all pulse durations tested). Likewise, the mean (± s.e.m) indirect activity rose by 25 ± 4.0 % when lengthening the stimulation time from 10 to 100 ms (average of all irradiances tested).

**Figure 4.**
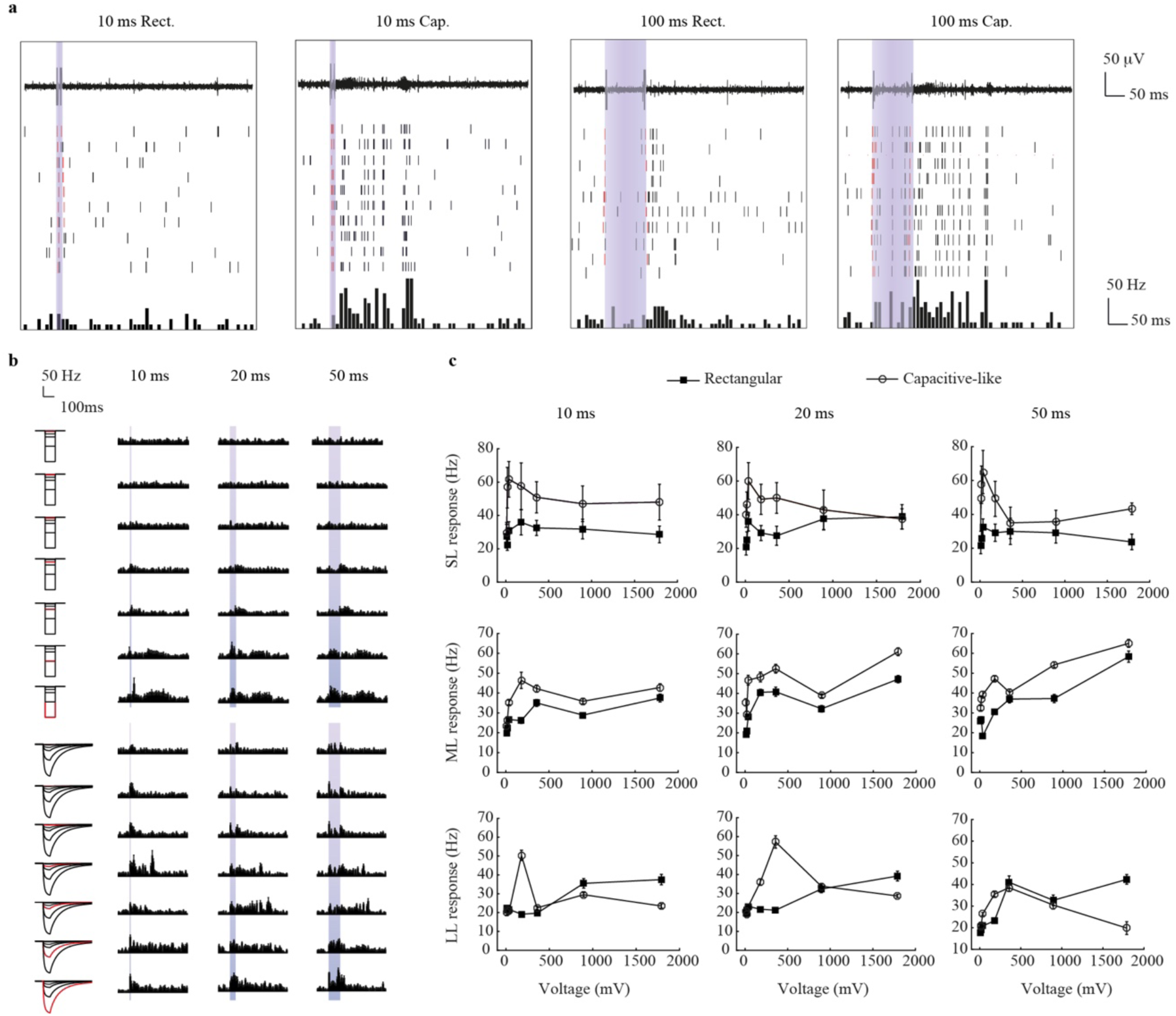
Comparison of RGCs response to rectangular and capacitive-like electric pulses. **a** Representative extracellular recordings of a single RGC in response to a 179 mV electrical stimulation with a 10 ms rectangular pulse (left), a 10 ms capacitive-like pulse (middle left), a 100 ms rectangular pulse (middle right), and a 100 ms capacitive-like pulse (right). The top row shows the electrophysiological recordings upon electrical stimulation; the middle row the raster plot of 10 consecutive sweeps; the bottom row shows the PSTHs (bin size 10 ms). The red raster lines correspond to the detection of the stimulus artefacts. **b** Mean (± s.e.m) PSTHs of RGC activity upon 10, 20, and 50 ms stimulation with rectangular (top) and capacitive-like (bottom) pulses (*n* = 13, for each RGC 10 sweeps have been averaged). The voltage pulses have been delivered with amplitudes of 8.95, 17.9, 35.8, 179, 368, 895 and 1790 mV. In **a** and **b**, the violet bar corresponds to the duration of the electric pulse. **c** Quantification of the mean (± s.e.m) firing rate of SL, ML and LL spikes upon electrical stimulation with 10 ms (left), 50 ms (middle), and 100ms (right) rectangular or capacitive-like pulses (*n* = 13, for each RGC 10 sweeps have been averaged).

### 3.3 Computational model

To provide a valid interpretation of the biophysical origin of the network-mediated spikes and its intricacy with the stimulus features, we simulated the response of stratified retinal tissue under voltage driven epiretinal stimulation. Four retinal cells layers have been modelled as a three-dimensional network (**Fig. 5a**): RGCs, amacrine cells (ACs), BCs, and horizontal cells (HCs). The capacitive-like voltage pulse has been modelled with a finite element analysis simulation as a local gaussian-shaped voltage increase, whose amplitude decreases with the depth within the retina (**Fig. 5d**). Given the high electrical resistivity of the neural retina, less than 5 % of the electric field reaches the PR layer when stimulating in epiretinal configuration, while only 31 % reaches the BC layer. Considering both the resistivity and the high capacitive properties of the neural retina, each layer has been modelled as a low-pass filter whose impulse response varies according to its distance from the stimulating electrode (**Fig. 5b**). The in-silico stimulation of the retina leads to a typical spiking pattern including SL and ML activities. Due to both the layered organization of the retinal cell types and the different low-pass filter they apply to the stimulus input, each layer reaches an activation peak in an asynchronous manner, as it can be seen from the simulated membrane potential of an HC, BC, AC, and RGC located above the centre of the stimulating electrode during a 20-ms rectangular stimulation (**Fig. 5c**). Direct voltage injection to the RGC layer leads to an action potential initiation within a few milliseconds after the stimulus onset (SL). However, since the stimulation does not have single cell resolution, the same stimulus propagates to the neighbouring retinal columns, including the close inhibitory surround. As a result, the excitability of the directly stimulated RGCs is decreased by their surrounding ACs. Meanwhile, granted that the input voltage is sufficient, upstream BCs are also depolarized and synaptically activating downstream RGCs. Voltage loss and filtering together engender a slow trade-off phase between excitatory and inhibitory backward inputs. The secondary activity latency is reproducible over stimulation conditions, as it depends on the connection balance between excitation and inhibition, that is to say, on the network itself, and not on the stimulation parameters. The burst of RGC secondary activity (ML) is voltage- and amplitude-dependent: it appears above a flux threshold of 2.0 µV s (**Fig. 5e**). Both BCs and ACs are necessary to produce a secondary activity pattern with a latency in the order of hundredths of a second (**Fig. 6**).

**Figure 5.**
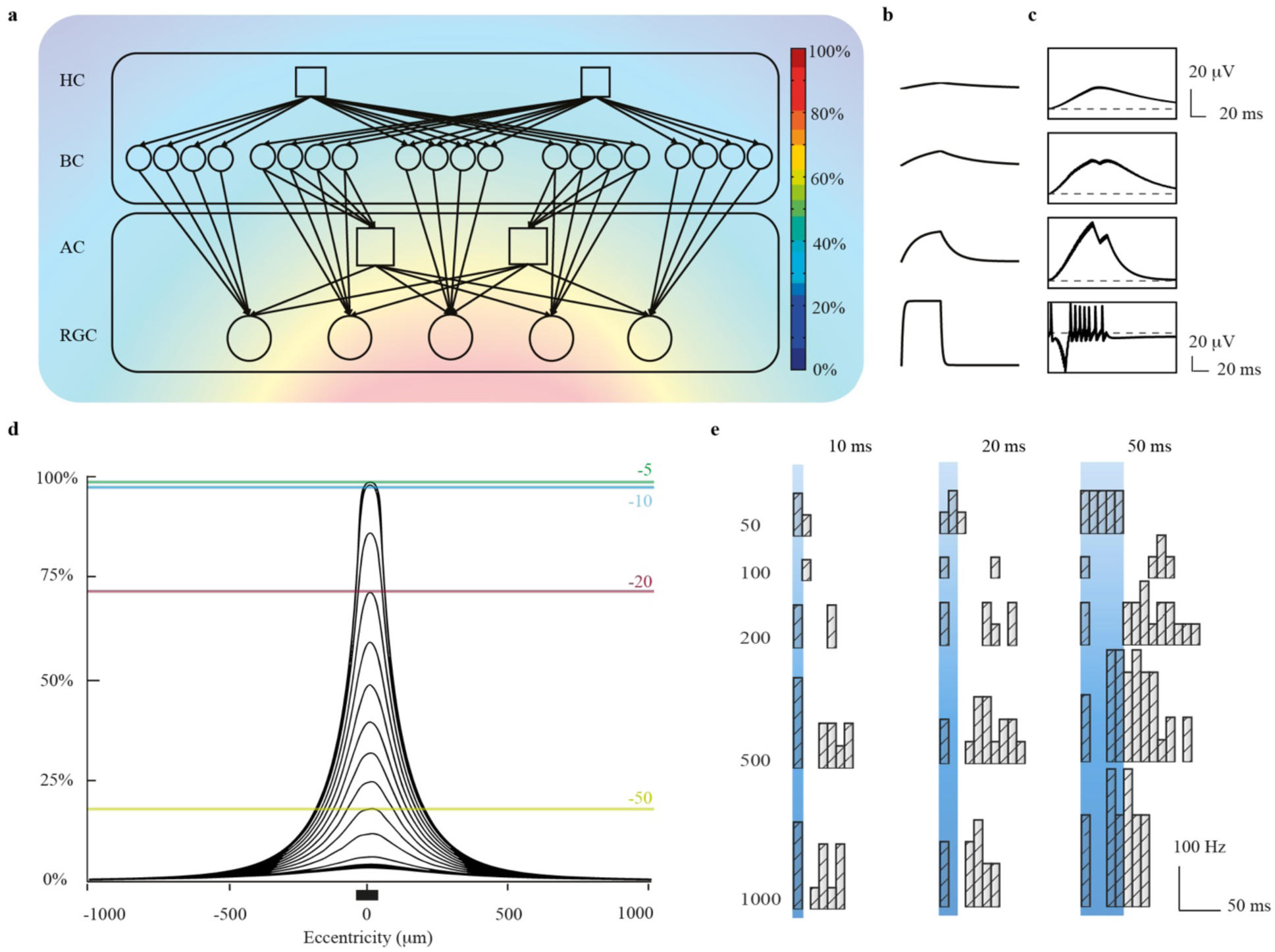
Biophysical model of the retinal layers under epiretinal stimulation. **a** Topology of the model. Circles represent the excitatory cells (RGCs, BCs), while squares the inhibitory cells (HCs, ACs). The background image shows a FEA simulation of the potential generated by an electrode placed in the centre of the sketched network. **b** Normalized impulse responses of HC, BC, AC, and RGC layers. **c** Membrane potentials of HCs, BCs, ACs, and RGCs located at the centre of the epiretinal electrode, upon a 50 ms / 180 mV rectangular stimulation. **d** FEA simulation of the potential generated at the photovoltaic electrode when it is facing the retinal tissue. The electrode is represented as a black bar. Each potential trace is separated from the previous one by a 5-µm step. The green line shows the potential at 5-µm depth from the electrode surface, the blue line at 10-µm depth, the red line at 20-µm depth, and the yellow line at 50-µm depth. **e** PSTH of a RGC located at the centre of the epiretinal electrode, upon 10, 20, and 50 ms stimulation. Rectangular voltage pulses have been delivered at 50, 100, 200, 500, and 1000 mV.

**Figure 6.**
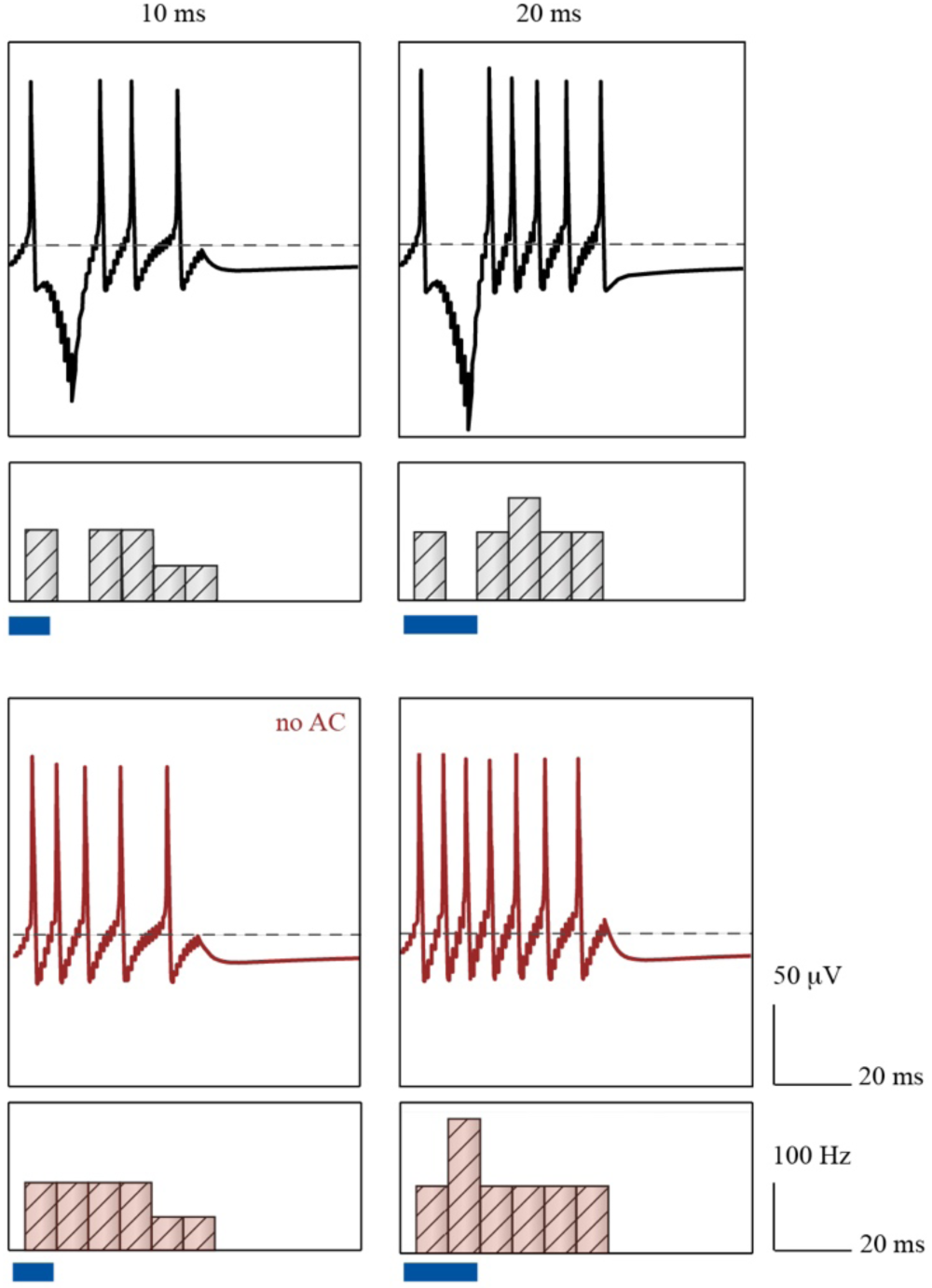
Modelling of RGC activity in the absence of ACs. PSTHs and membrane potentials of the RGC located at the centre of the electrode, upon 10 and 20-ms capacitive-like stimulation at 250 mV, with (black) and without (red) presynaptic inhibition from the ACs.

We then stimulated the biophysical network with either rectangular or capacitive-like voltage pulses. The increasing and decreasing phases of capacitive-like voltage pulse have been here fitted as one-term exponentials. Membrane potentials’ rise in ACs, BCs, and to a lower extent in HCs is observed to be slower but of higher magnitude when the stimulus is capacitive-like shaped with respect to the rectangularly shaped one (**Fig. 7a**). Rapid voltage transitions also generate fast interneurons membrane potential rise, but without any sustained potential. As a consequence, capacitive-like pulses generate in RGCs an average (± s.e.m) indirect firing activity of 52.6 ± 20.0 Hz, while rectangular pulses generate an average (± s.e.m) indirect firing activity of 37.1 ± 12.3 Hz (**Fig. 7b**). For identical over-threshold conditions eliciting non-zero indirect activity with both pulses’ shapes, capacitive-like pulses generate an average (± s.e.m) indirect activity 66.6 ± 11.7 % higher than the one obtained with rectangular pulses. Direct activation of RGCs in-silico is not affected by the stimulus shape.

**Figure 7.**
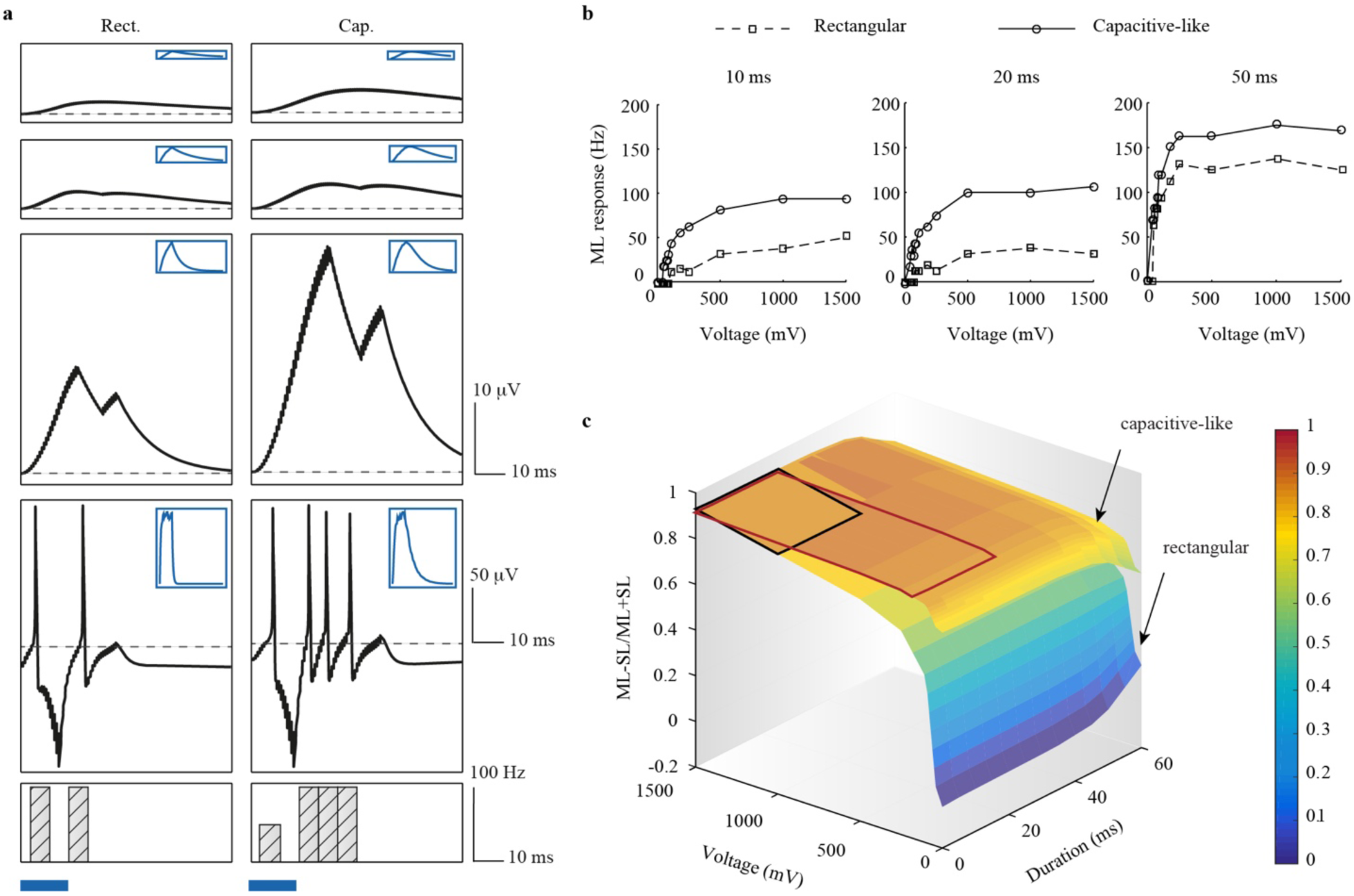
Comparison of the retinal circuit response upon stimulation with rectangular and capacitive-like voltage pulses. **a** PSTHs and membrane potentials of HCs, BCs, ACs, and RGCs located at the centre of the epiretinal electrode, upon 180 mV / 20 ms, square (left) and prosthetic (right) stimulations. The top right corner boxes show the normalized impulse response for each cell layer. **b** Indirect (ML) spiking activity generated in the retinal ganglion cell located at the centre of the electrode by 10, 20, and 50 ms stimuli for both voltage shapes. Delivered peak voltages ranged from 40 to 1500 mV. **c** Parametric model of the computationally predicted RGC activity. The indirect over direct activity ratio has been modelled as a function of the delivered pulse duration and voltage. The red and black squares respectively highlight the optimal parameters found for capacitive-like and rectangular pulses (duration < 20 ms, ML-SL/ML+SL > 0.8).

Recent evidence pointed out the challenge of stimulating from the epiretinal side the deep ganglion cell layer without promoting local hyperpolarization of axons of passage [25, 26]. Epiretinal indirect stimulation of RGCs is, therefore, a promising strategy to overcome the problem of axonal depolarization and the resulting loss of resolution. In this perspective, the use of non-rectangular pulses can successfully shift the RGC activation pattern from direct to indirect activation. **Fig. 7c** shows a linear assumption of the indirect-to-direct firing rates ratio for various pulse voltages and durations, either rectangularly or capacitive-like shaped. While rectangular stimuli require high voltages (higher than 1 V) or alternatively very long durations (longer than 60 ms) to maximize the indirect activity with respect to the direct one, similar activity ratios can be obtained with capacitive-like pulses at voltages one order of magnitude lower. Symmetrically, capacitive-like pulses allow maximising the indirect-to-direct activity ratios with shorter pulses compared to those necessary with rectangular pulses, which would be hardly compatible with high-frequency stimulation. Optimal parameters for rectangular and capacitive-like pulses are highlighted respectively in black and red (**Fig. 7c**).

In addition, since capacitive-like pulses depolarize qualitatively longer the inner retinal cells and especially BCs, it facilitates the temporal summation of repetitive stimuli (**Fig. 8**). The conventional stimulation frequencies used in retinal prostheses range from 5 to 20 Hz [8,9,24,27,28], and recent evidence appoint 10 Hz as an optimal stimulation frequency for epiretinal prostheses within this range [43]. The trade-off between stimulation frequency and stimulus duration becomes especially important when dealing with photovoltaic electrodes. A compromise has to be found between maintaining a pulse short enough to avoid photothermal damage to the retina and still enabling indirect activity and eventually temporal summation. Mastering the decay speed of the voltage stimulus can be one approach to do so. Tuning the discharge capacitive properties of the electrode/electrolyte interface can modulate the ML activity elicited by a pulse of identical duration and voltage (**Fig. 9**). For similar over-threshold single 10-ms pulses, doubling the decay phase time constant increases the mean (± s.e.m.) ML activity of 39 ± 12 % (**Fig. 9b,c**). No comparable modulation could be found when varying the rising phase time constant. Last, shuffling the stimulating electrode from epiretinal to subretinal side allowed to shorten the onset to indirect activity burst delay (**Fig. 10**), as previously reported in-vitro [31].

**Figure 8.**
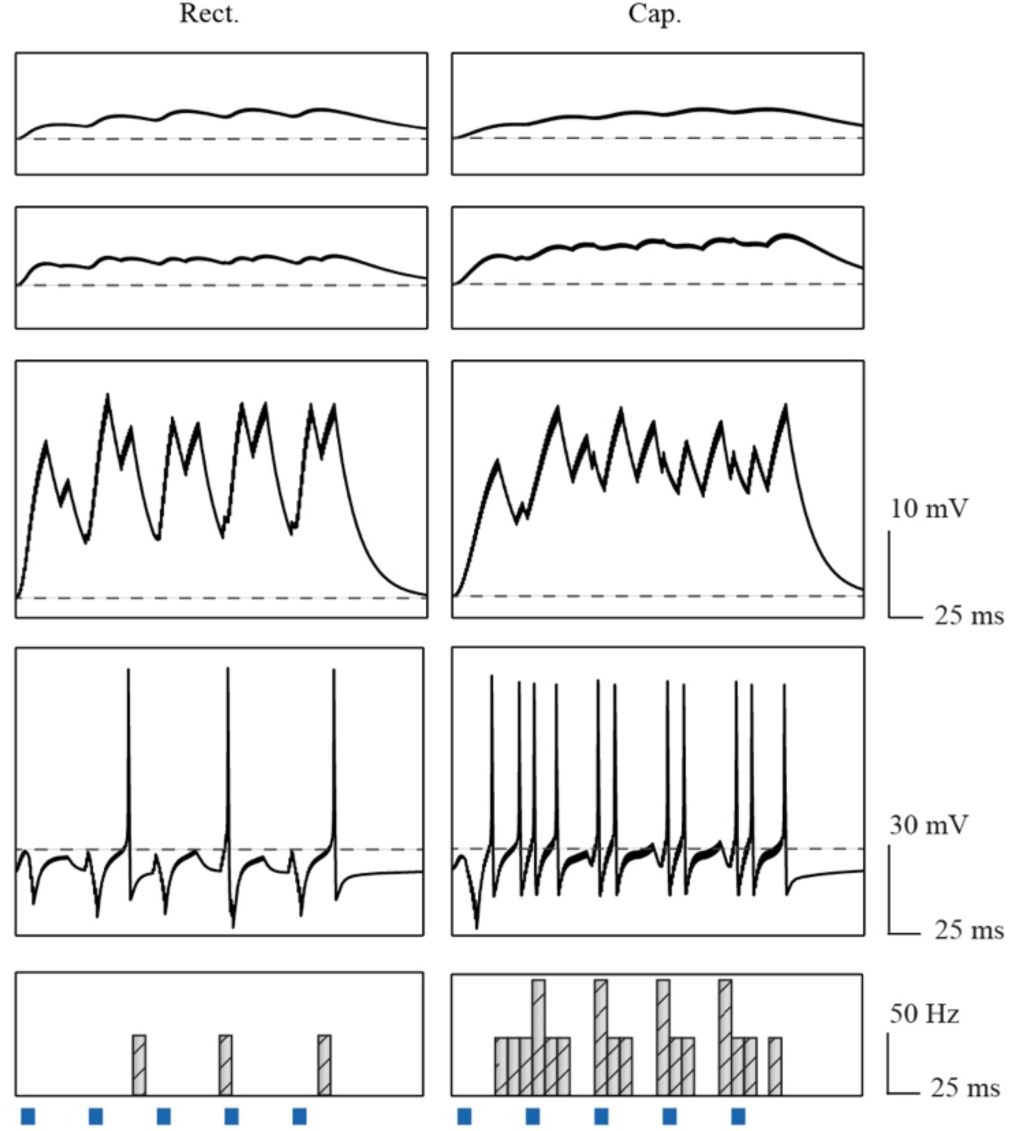
Modelling of the retinal network activity upon repetitive stimulation. PSTHs and membrane potential of HCs, BPc, ACs, and RGCs located at the centre of the electrode, upon repetitive rectangular (left) and capacitive-like (right) stimulation. Voltage pulses of 10 ms at 20 mV pulses have been delivered at 20Hz.

**Figure 9.**
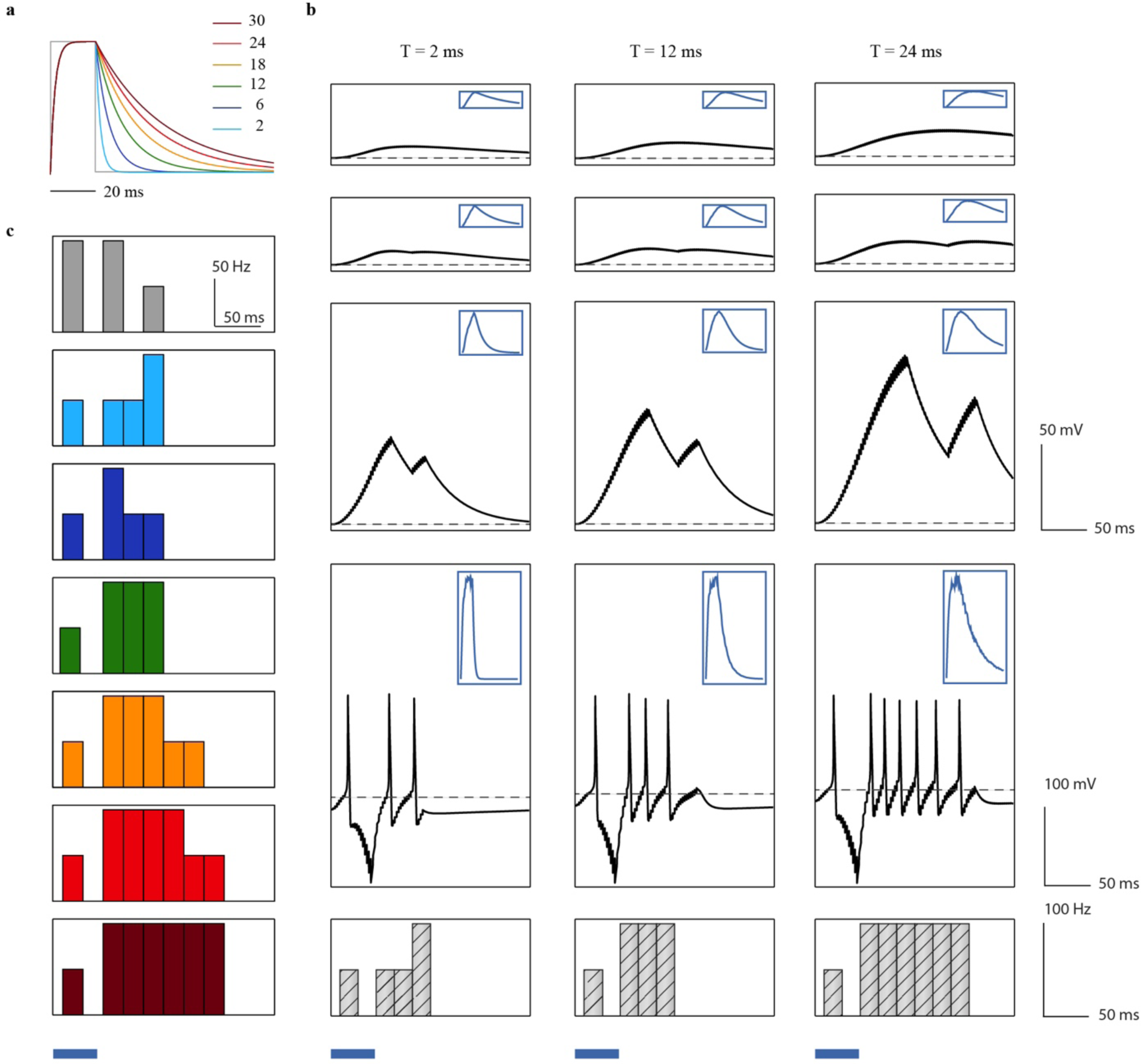
Comparison of network-mediated activity elicited by various capacitive-like stimulus shapes. **a** Theoretical normalized photovoltage curves with various discharge time constants. The photovoltage measured on top of photovoltaic pixels[15] is fitted by the green curve. **b** PSTHs and membrane potentials of HCs, BPs, ACs, and RGCs located at the centre of the electrode, upon 20 ms capacitive-like stimulation at 250 mV, with discharge time constants of 2, 12, and 24 ms. The top right corner boxes show the normalized impulse response for each cell layer. **c** PSTH of the RGC located at the centre of the electrode, upon 20 ms rectangular and capacitive-like stimulations at 250 mV. Discharge time constant of the capacitive-like stimuli was set to 2 (light blue), 6 (blue), 12 (green), 18 (orange), 24 (red), and 30 (vinous) ms.

**Figure 10.**
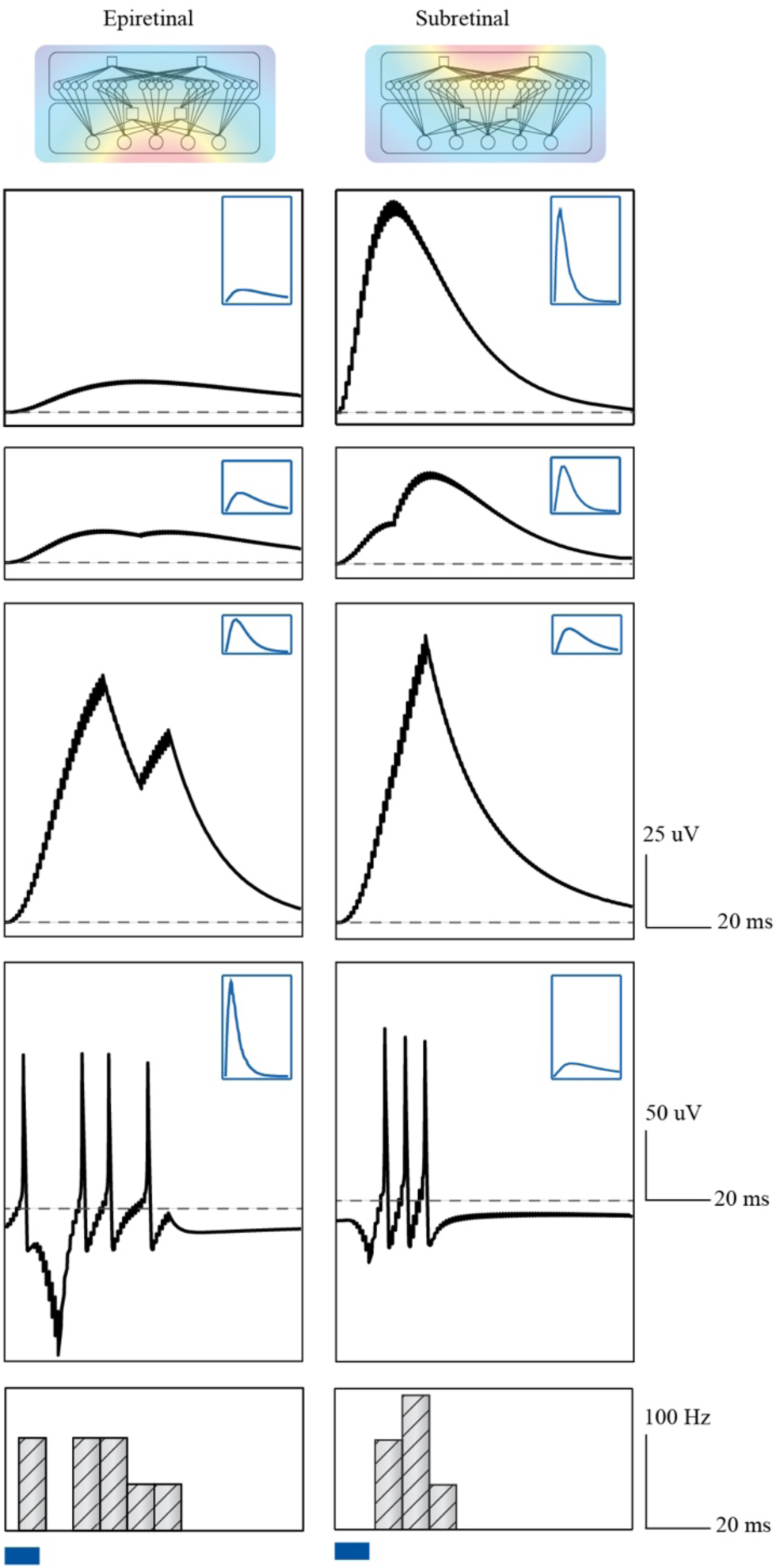
Comparison of epiretinal and subretinal stimulation in-silico. PSTH and membrane potential of HCs, BCc, ACs, and RGCs located at the centre of the electrode, upon 10-ms capacitive-like stimulation at 250 mV from an epiretinal or a subretinal electrode. Top right corner boxes show the normalized impulse response for each cell layer. For subretinal stimulation, the electrode was placed 10 µm away from the horizontal cell layer.

### 3.4 Spatial selectivity of rectangular and capacitive-like voltage pulses

Ultimately, we addressed the spatial selectivity of capacitive-like voltage pulses using a computational/experimental hybrid approach. From two-dimensional activation plots of simulated RGC, AC, BC and HC populations, we estimated the spatial extent of the layers’ response to electrical stimulation for rectangular and capacitive-like, short and long pulses, below or above the indirect activation threshold. **Fig. 11** shows the inverse relationship between the spatial extent of the AC layer activation and the spatial extent of the RGC spiking response to the stimulation. For similar voltage peaks and pulse durations, capacitive-like pulses can elicit not only higher membrane potential changes in the inhibitory interneurons than rectangular pulses, but also affect a wider pool of them, as a result of the gaussian-shaped voltage probability distribution. This strengthening and widening of the AC response are again observed to be voltage, duration and shape-dependent (**Fig. 11a,b**). The wider activation of the recorded AC pool is observed under capacitive-like 50-ms over-threshold stimulation, in which a 7.5 cells-diameter area is activated around the electrode. Though, while lengthening the stimulation pulse, the gap between AC activation profiles triggered by one or the other pulse shapes reduces, presumably due to the saturation of network response kinetics. Parallelly, the spatial extent of RGC indirect activity is also observed to be voltage, duration and shape-dependent. Above the indirect activation threshold, the AC activation spread and the RGC indirect response spread (estimated from a Gaussian fit of the membrane potentials of the 10 × 10 cell population) can be linearly anticorrelated (**Fig. 11c**).

**Figure 11.**
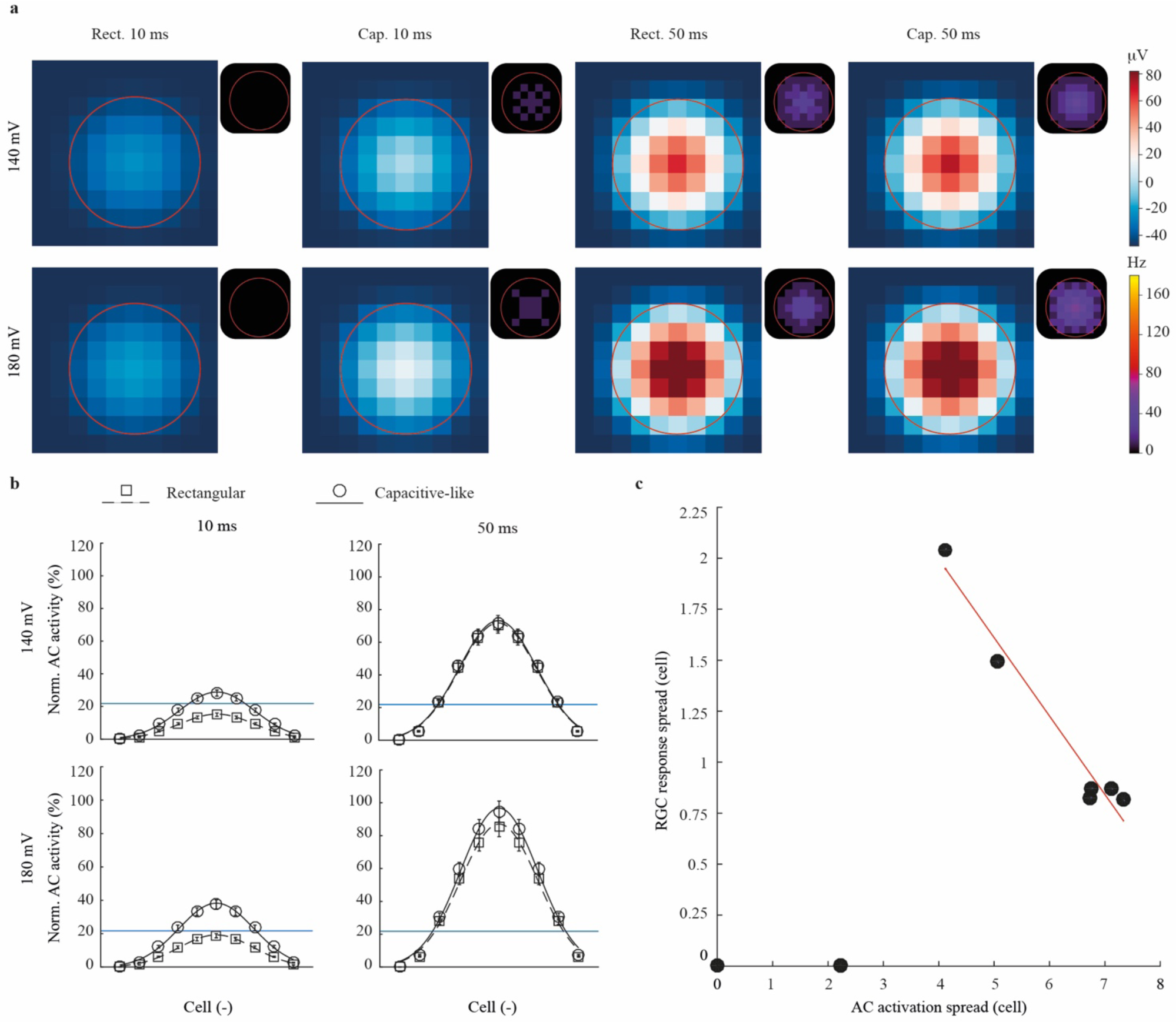
Spatial modelling of the retinal circuit response. **a** Colour map of AC membrane potential rising peak. A 10 × 10 pool of ACs located around the stimulation electrode has been recorded upon short (10 ms) and long (50 ms) rectangular or capacitive-like voltage pulses at 140 mV (top) and 180 mV (bottom). Each pixel represents the cell membrane potential difference with respect to the membrane potential threshold required for the appearance of the ML activity. Positive differences are represented in white to red colours, while negative (under-threshold) differences are represented in blue. The RGC indirect activity (ML) corresponding to each condition is plotted on the top right corner of each colour map. The red circle indicates the electrode location. **b** Mean normalized activation profile and Gaussian fit of the activity of inhibitory (AC) cells upon 10 and 50 ms rectangular or capacitive-like voltage pulses peaking at 140 mV (top) and 180 mV (bottom). 10 × 10 cells have been averaged over 4 directions. The blue line indicates the membrane potential threshold for indirect activity. **c** Spatial extent of RGCs indirect response with respect to the extent of AC activity. The spatial extent of the RGC response has been calculated as the full width at half maximum of its activation profile fit, while the spatial extent of the AC activation has been calculated as the full width at the threshold for indirect activity.

We then estimated the eRF of stimulated RGCs in-vitro. Rd10 retinas have been explanted on the custom 80-µm titanium MEAs and RGCs have been located by recording extracellularly their spontaneous activity. Each RGC has been successively stimulated from the 20 nearest electrodes with rectangular and capacitive-like short and long pulses. The portion of the MEA activating either directly or indirectly the recorded cell is labelled as SL-eRF, ML-eRF, or LL-eRF. eRFs for long, short, capacitive-like, and rectangular pulses are shown in **Fig. 12**. Similar to previous experiments, capacitive-like stimuli delivered through the closest (central) electrode could elicit stronger activity than rectangular stimuli. However, not only the closest electrode has been able to activate the targeted RGCs. SL-eRFS notably present elongated shapes (**Fig. 12c**), presumably due to axonal stimulation. An action potential generated in a distal axonal segment and antidromically propagated in less than 10 ms, would indeed be classified as direct RGC activation (see Methods).

**Figure 12.**
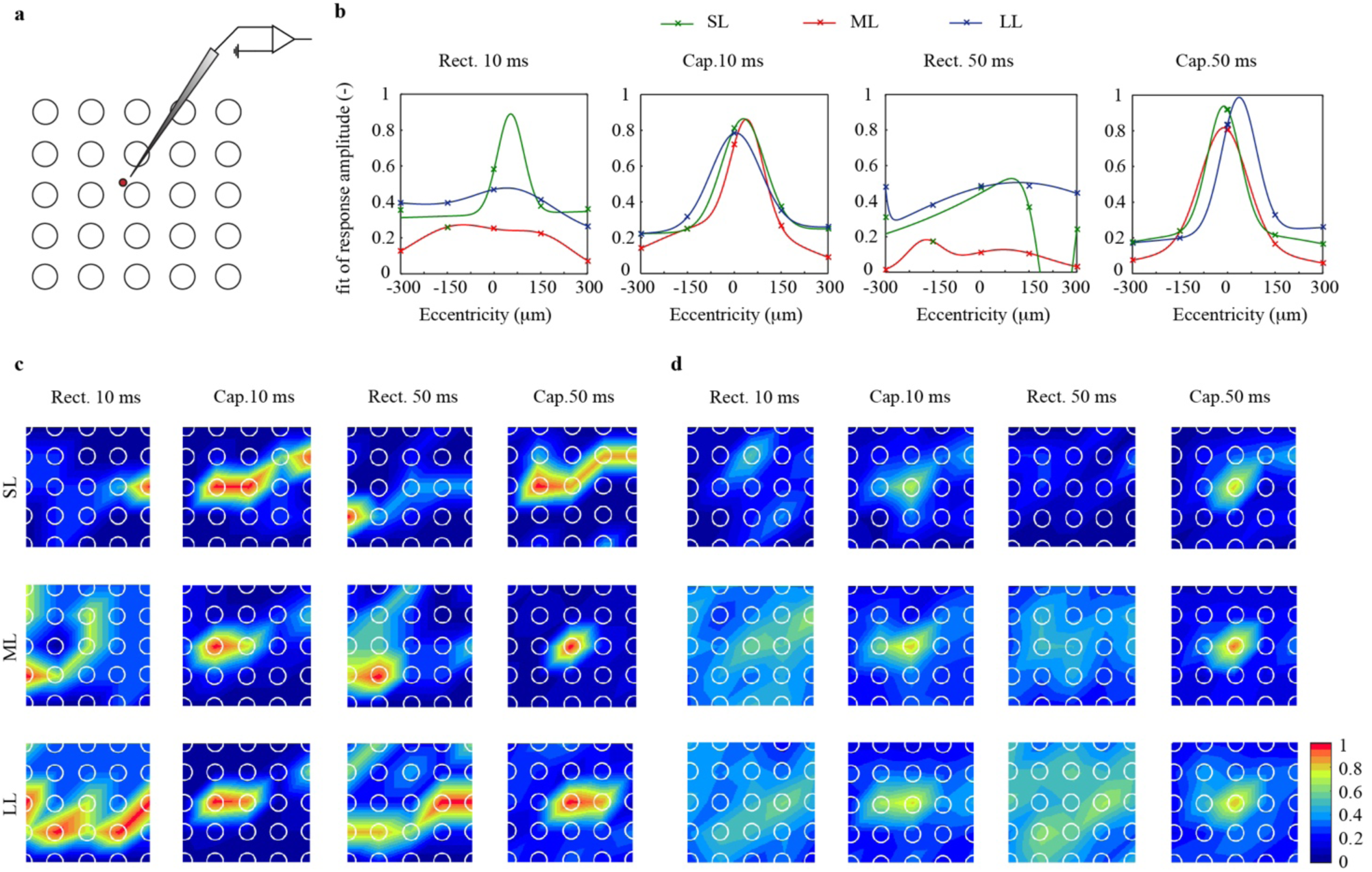
Electrical receptive fields of RGC upon rectangular and capacitive-like stimulation. **a** Sketch of the stimulating and recording electrodes. The approximated soma location of the recorded cell is highlighted in red. **b** Fit of the grand average SL (green), ML (red), and LL (blue) responses amplitudes. Experimental measures obtained with individual electrodes stimulation have been averaged over horizontal, diagonal, and vertical axis and fitted with a Gaussian function (*n* = 8, for each RGC 10 sweeps have been averaged). eRF diameters have been calculated as the minimal distance corresponding to the 3^rd^ quartile of the activity distribution. SL, ML, and LL eRF diameters have been quantified in 101, 107, and 218 µm with short rectangular stimulation; 226, 111, and 220 µm with long rectangular stimulation; 82, 93, and 110 µm with short capacitive-like stimulation; and 66, 76, 86 µm with long capacitive-like stimulation. **c** Representative heatmap of normalized SL, ML, and LL activities recorded with short (10 ms) and long (50 ms) rectangular or capacitive-like, both with peak voltages of 180 mV (*n* = 1, 10 sweeps have been averaged). **d** Mean heatmap of normalized SL, ML, and LL activities recorded with short (10 ms) and long (50 ms) rectangular or capacitive-like pulses with peak voltages of 180 mV (*n* = 8, for each RGC 10 sweeps have been averaged). **c** and **d** heatmaps were generated from linear interpolation of experimental SL, ML, and LL values recorded from individual electrode stimulations.

Regarding indirect eRFs, the stimulation conditions providing the most focused response are the ones previously associated with a high indirect activity, namely short and long capacitive-like voltage pulses. Besides the weak values of mean network-mediated activity (due to idiosyncratic preferred stimulation axis), rectangular pulses exhibit broad indirect eRFs (**Fig. 12b,c**). Rectangular ML-eRFs and LL-eRFs had respective diameters of 147 and 218 µm (fit of *n* = 8 cells from *N* = 8 retinas, pulse duration 10 ms and amplitude 179 mV). On the contrary, capacitive-like pulses exhibit ML-eRFs and LL-ERFs clustered around the central electrode, with respective diameters of 93 and 110 µm (fit of *n* = 8 cells from *N* = 8 retinas, pulse duration 10 ms and amplitude 179 mV). Furthermore, the longer the pulse duration, the more clustered indirect eRFs is observed. The narrower eRFs (ML-eRF diameter = 76 µm, LL-eRF diameter = 86 µm) has been observed for capacitive-like stimulation of 50 ms.

## 4. Discussion

In this work, we used the POLYRETINA photovoltaic prosthesis [15], whose 80-µm pixels generate cathodic capacitive-like voltage pulses under green light stimulation, to generate network-mediated activation of RGCs from dystrophic light-insensitive retinas. In our experiments we have observed that the epiretinal photovoltaic stimulation of RGCs generates indirect responses 30 % (ML) and 8 % (LL) higher than indirect responses elicited by electrical stimulation with rectangular pulses under similar conditions (pulse duration and peak amplitude).

Overall, indirect network-mediated activity (ML and LL) could be enhanced by irradiance or voltage increase, pulse lengthening, and substitution of rectangular pulses into non-rectangular capacitive-like pulses, all resulting in larger activation of inner retinal interneurons. Both gradual voltage decrease and sustained voltage delivery showed the ability to maintain non-spiking BCs and ACs active for tenths of seconds after the stimulus onset, leading to the characteristic RGC activation pattern consisting in a direct single spike from local membrane depolarization, an inhibition period, and a secondary wave of activation. The contribution of the inhibitory network to this pattern allows a clustered RGC indirect activation, compared to the spatial extent of the direct response.

We have demonstrated that our photovoltaic approach, despite the apparent loss of temporal precision due to the capacitive-like voltage transients, is efficient to trigger realistic network-mediated activity in blind retinas upon epiretinal stimulation. The relevance of the voltage pulse shape and the charge delivery rate is in line with recent reports about the efficiency of non-rectangular pulses to trigger network-mediated activity [42] and the sensitivity to low-frequency domains of inner retinal cells [43]. Fast responding spike-encoding RGCs show preferential sensitivity to rectangular stimuli, while the activation of voltage-encoding interneurons can be modulated by slower voltage changes such as capacitive-like photovoltaic pulses. However, the biophysical explanation for this frequency shift remains to be investigated. The indirect activity delay shrinkage that we observe when displacing the electrode towards the subretinal space (**Fig. 10**), as it was previously observed in-vitro [31], suggests that the input filters are not electrophysiological intrinsic properties of the cell types *per se*, but rather consequences of the layered structure of the retina and its electrical resistivity. Reports of direct response within a timescale of several milliseconds in subretinal configuration, namely twice as long as delays reported in epiretinal configuration direct stimulation, also supports this hypothesis [31, 51].

The activation of axon bundles keeps being one of the most challenging aspects of epiretinal stimulation. Our previous work with POLYRETINA revealed a direct firing probability of about 34 %, and we reported in the present study a mean (± s.e.m) direct firing activity of 38.9 ± 1.2 Hz. However, in the perspective of axons stimulation avoidance, and given the ability of the photovoltaic approach to efficiently trigger indirect activation, it seems more and more realistic to design a stimulation protocol for epiretinal indirect stimulation thanks to capacitive-like waveforms and long pulse durations (longer than 20 ms).

In addition to medium latency spikes, a third indirect and further delayed RGC burst of activity (LL) was reported in this work and others [31, 38] at high stimulation voltages. The identification of sustained RGCs as generators of successive oscillatory bursts under electrical stimulation, together with the absence of LL activity in-silico suggests that they might have a functional type related origin. It remains still unclear how the successive waves of activation would be further processed and interpreted by the visual cortices. The perception of elongated ovoid phosphenes is imputed to the bidirectional depolarization of axonal fibres [25, 26], that is to say, the temporal precision separating orthodromic and antidromic followed by orthodromic signals vanished with upstream signal integration. However, indirect waves with a delay in the range of tenths of seconds may be more prone to reproduce the natural temporal structure of visual responses. Indeed, though many individual spikes can be timed with millisecond precisions to the visual stimulus they encode, the relative precision among the activated RGC population is a higher critique feature for information integration in the thalamus [52]. Approaching the relative temporal structure of natural vision could provide an equally accurate representation of the slowly changing visual word. The ability of non-rectangular pulses to generate a sustained membrane potential rise in retinal interneurons also facilitate the temporal summation of under-threshold stimuli (**Fig. 8**), making it conceptually feasible to abolish any direct depolarization of nearby RGC axon initial or distal segment, while promoting indirect summed activity.

Turning epiretinal stimulation into an explicitly indirect strategy is also promising for prosthetic response resolution. Our results suggest a key role of ACs in the generation of a network-mediated temporal pattern of activity. In addition, recruiting the natural lateral inhibition circuit allows to spatially cluster the eRFs of targeted RGCs. Indeed, the assessment of RGCs’ eRFs reveals disparities between the spatial distribution of direct and indirect excitabilities. The clustering of the RGC eRF can notably be impaired by several factors; the first of them being the local depolarization of the RGC axon by a peripheral electrode, as observed in the direct individual eRFs and further cancelled out at the population level. The spatial resolution of the stimulation itself is another spreading factor, considering that a single 80-µm diameter electrode can be estimated to cover the natural RF of 15 to 20 RGCs in an intact mouse retina, but up to 160 RFs in the human fovea [53, 54]; and that the extra potential generated around the electrode can be estimated to cover 10 additional RFs. Furthermore, gap junctions between RGCs and eventually aberrant connection with peers or other miswired partners could alter the spatial clustering of the individual RGCs responses. Those three latest points can be attenuated by decoupling the RGC eRFs ones from the others thanks to their respective inhibitory surrounds. The sustained stimulation of the inner retina and the lateral inhibitory network, via long pulses or non-rectangular waveforms, contribute to the clustering of the indirect eRF.

The improvement of the response resolution can be achieved with non-rectangular pulses, but also with long pulses (longer than 20 ms). Indeed, both strategies exploit the same interneuron’s continuous electrical properties. Various evidence, including ours, demonstrates that long pulses elicit strong network-mediated activity in RGCs. Our computational-experimental approach suggests that the RGC network-mediated activity is necessarily associated with a partial activation of the AC layer and backwards inhibition of the eRF surround. This clarifies the response resolution refinement obtained in-vitro and in-vivo with pulses exceeding 25 ms [24]. In the above-mentioned study, the calcium imaging readout of spatial RGC activation in-vitro could be clearly narrowed by the lengthening of electrical stimulation, as complementary do the indirect eRFs. Similarly, the indirect eRFs clustering that we observed with capacitive-like pulses, that exploits the same mechanisms, is expected to narrow the spatial extent of RGC layer activation, as computationally simulated. However, pulses duration dramatically limits the affordable stimulation frequency: an optimal duration pulse of 25 ms would theoretically limit the prosthesis operating range to 20 Hz (or less if we consider a safe interval between consecutive pulses), far below flicker fusion threshold. Non-rectangular waveforms such as our capacitive-like stimuli allow reaching indirect firing rates comparable to those obtained with 20-ms rectangular pulses with pulses two times shorter, thus increasing the theoretical stimulation rate limit. Along with this, for identical pulses duration, the use of non-rectangular pulses enables to lower the voltage threshold necessary to mobilize both excitatory and inhibitory inner retinal cells.

Hereabout, the direct as the indirect activation of rd10 RGCs is significantly higher when the explants have been stimulated with capacitive-like pulses compared to rectangular stimuli. We classified spikes as direct (SL) when they happen to occur within the first 10 ms; nevertheless, due to the stimulation artefacts, spikes occurring within 1.5 ms after the stimulus onset could not be detected. Because slower charge increase delays the direct spikes timing by up to 2 ms [42], the amount of undetected direct spikes minors with a capacitive-like stimulus.

## 5. Conclusion

Sustained activity of inner retinal neurons can be achieved by delivering non-rectangular voltage pulses and/or by lengthening the pulses’ duration. Such stimulation paradigm allows to indirectly target RGC in a realistic and focused approach, while benefiting from the implantation convenience of epiretinal devices.

Optimal pulse shape engineering is a complementary strategy to further promote network-mediated response without lengthening the stimulation pulses. A slower discharge rate of the electrode can notably sustain network excitation and increase resulting RGCs firing rate up to three times the rate recorded with comparable rectangular pulses. Moreover, our results demonstrate that recruiting inner retina cells with epiretinal stimulation enables not only to bypass axonal stimulation but also to obtain a more focal activation thanks to the natural lateral inhibition. Additional in-vitro evidence suggests a conjoint role of the ascending stimulus ramp on the ability to generate indirect activity [42].

In this perspective, the use of capacitive-like waveforms generated by photovoltaic prostheses (e.g. POLYRETINA) may allow improving the neural response resolution while standing high-frequency stimulation. At last, the relevance of the strategies for network-mediated activity enhancing to improve artificial vision resolution will require to be established by evaluating the restored visual acuity both in in-vitro and in-vivo.

## Acknowledgements

This work has been supported by École Polytechnique Fédérale de Lausanne, Medtronic, and Fondation Pierre Mercier pour la science. We would like to thank Alban Bornet and Anton Voronov for their help in Python.

## Author contributions

N.A.L.C. designed the study, performed all the experiments and simulations, and wrote the manuscript. M.J.I.A.L. fabricated the photovoltaic array and the custom MEA for electrical retinal stimulation and designed FEA simulations. D.G. designed and led the entire study. All the authors read, edited and accepted the manuscript.

## Competing Financial Interests statement

The authors declare no competing financial interests. Correspondence and requests for materials should be addressed to D.G.

